# The Amygdalostriatal Transition Area Exhibits Lateral Amygdala–Like Spiking Activity and Tone–Shock Pairing–Induced Plasticity

**DOI:** 10.64898/2026.06.12.731865

**Authors:** Dániel Magyar, Mária Rita Karlócai, Norbert Hájos

## Abstract

During Pavlovian fear conditioning, presentation of a conditioned stimulus, such as a tone, together with an unconditioned stimulus, such as an electrical shock, excites neurons in the lateral amygdala (LA). Prevailing models propose that sensory stimulus-evoked activity in the LA is subsequently propagated to its downstream nuclei: the basal amygdala (BA) and central amygdala (CeA). To test this assumption, we performed in vivo electrophysiological recordings in awake, head-fixed male and female mice. We found that tone presentation did not elicit a significant increase in firing in BA or CeA neurons. In contrast, shock presentation evoked similarly robust spiking responses in LA and BA neurons but only a modest increase in CeA neurons. Notably, neurons in the amygdalostriatal transition area (AStria) exhibited LA-like sensory stimulus-evoked responses at both short (<25 ms) and longer (<500 ms) timescales. To examine the role of feedforward inhibition in tone- and shock-evoked activity, we investigated the contribution of parvalbumin interneurons using optogenetics and found that short-latency (<25 ms) spiking in both the LA and AStria was regulated by these inhibitory cells. Finally, LA and AStria neurons exhibited remarkably similar response types, spiking dynamics, and pairing-induced plasticity during repeated tone presentations, subsequent tone–shock pairings, and post-pairing tone presentations. Together, these findings support a model in which the LA and AStria operate in parallel, similarly integrating tone and shock signals during fear conditioning, whereas BA and CeA neurons are not robustly recruited by these sensory stimuli under the conditions tested.

## Introduction

The formation of long-term fear memories is essential for survival and is mediated, at least in part, by amygdala circuitry (Fendt and Fanselow, 1999; LeDoux, 2000; Maren, 2001). Prior work has identified the lateral amygdala (LA) as a critical site where, during Pavlovian fear conditioning, a neutral conditioned stimulus (CS; e.g., a tone) and an aversive unconditioned stimulus (US; e.g., a footshock) converge onto a small subset of neurons, resulting in the formation of a fear memory engram (Bordi et al., 1993; Romanski et al., 1993; Quirk et al., 1995; Quirk et al., 1997; Collins and Pare, 2000; Pare and Collins, 2000; Goosens et al., 2003; Reijmers et al., 2007; Johansen et al., 2010; Rashid et al., 2016; Abdou et al., 2018; Abatis et al., 2024).

Auditory CS information reaches the LA via two parallel routes. Thalamic nuclei, including the medial part of the medial geniculate body and posterior intralaminar nucleus (MGm/PIL), elicit rapid spiking with latencies of ∼15–25 ms. In contrast, auditory cortex-driven firing in the LA occurs later, with at least one additional monosynaptic delay, resulting in an extra ∼8–10 ms latency (LeDoux et al., 1985; LeDoux et al., 1991; Quirk et al., 1995; Quirk et al., 1997; Collins and Pare, 2000; Repa et al., 2001; Goosens et al., 2003). Thus, tone-evoked responses with latencies <25 ms are driven by thalamic input. Although several sources of US input to the LA have been proposed (Shi and Davis, 1999; Lanuza et al., 2008; Johansen et al., 2010), a recent study showed that the MGm/PIL pathway can trigger footshock-evoked firing in the LA (Taylor et al., 2021), and that some LA-projecting thalamic neurons in these nuclei are activated by both tone and footshock (Ledoux et al., 1987; Bordi and LeDoux, 1994; Lanuza et al., 2008; Taylor et al., 2021). These findings raise the question of how sensory stimulus-driven excitation shapes LA activity and whether tone- and shock-evoked inputs interact when presented together.

In the prevailing model, CS–US-evoked spiking activity in the LA instructs downstream regions, including the basal amygdala (BA) and central amygdala (CeA), to generate appropriate defensive behaviors based on learned fear (Maren and Quirk, 2004; Johansen et al., 2011; Herry and Johansen, 2014; Tovote et al., 2015). However, this sequential model of information flow within the amygdala is difficult to reconcile with lesion studies suggesting that the LA and BA may play distinct roles in auditory-cued fear learning (Nader et al., 2001; Manassero et al., 2018); but see (Goosens and Maren, 2001). Distinct functions of LA and BA neurons are further supported by differences in their connectivity (Hintiryan et al., 2021; Reeb et al., 2025) and gene expression profiles (O’Leary et al., 2020; Hochgerner et al., 2023; Lim et al., 2024). Importantly, an in vitro study suggested that the LA may transmit information primarily to the neighboring amygdalostriatal transition area (AStria), rather than to the BA or CeA (Wang et al., 2002). Furthermore, the AStria shares thalamic and auditory cortical inputs with the LA (LeDoux et al., 1991; Romanski and LeDoux, 1993), suggesting that this striatal region may also participate in sensory processing and the regulation of learned fear. Indeed, recent studies have shown that AStria neurons contribute to fear-memory processes (Mills, 2022; Kintscher et al., 2023). Yet, how AStria neurons respond to sensory stimuli and how their spiking activity evolves during repeated and paired tone–shock presentations remain unclear.

To address these questions, we performed in vivo electrophysiological recordings in awake, head-fixed mice and compared sensory stimulus-evoked spiking across four neighboring amygdala regions. We found that LA and AStria neurons exhibited remarkably similar spiking activity and pairing-induced plasticity following tone–shock pairing. These findings may prompt a re-evaluation of prevailing models of information flow downstream of the LA, as the AStria, rather than the BA or CeA, may be the principal target recruited during the initial processing of tone and shock stimuli.

## Materials and Methods

### Animals and Surgical Procedures

All experimental procedures were performed in accordance with the National Institutes of Health Guide for the Care and Use of Laboratory Animals and were approved by the Indiana University Bloomington IACUC. A total of 41 mice (11 male and 2 female PV-Cre mice plus 15 male and 13 female wild-type mice) were used. Adult C57BL/6 mice (8–12 weeks; The Jackson Laboratory) or PV-Cre mice (RRID:IMSR_JAX:017320) were single-housed after surgery on a 12-h light/dark cycle with ad libitum food and water.

For awake head-fixed recordings, mice were anesthetized with isoflurane (1–2% in O_2_) and implanted with a custom headplate (Luigs & Neumann GmbH, Ratingen, Germany). For optogenetic experiments in PV-Cre mice, AAV5-CAG-FLEX-ArchT-tdTomato (gift from Edward Boyden, Addgene viral prep # 28305-AAV5; http://n2t.net/addgene:28305; RRID: Addgene_28305) was injected (100 nL per depth) into the right AStria (AP: −1.7, ML: +3.8, DV: −3.0 mm from bregma, at a 10° angle) and LA/BA (AP: −1.7, ML: +4.0, DV: −3.6 mm from bregma, at a 10° angle). An optic fiber (400 µm diameter; Thorlabs GmbH, Bergkirchen, Germany) was implanted (AP: −1.7, ML: +2.5, DV: −3.5 mm at a 10° angle). Implants were secured with dental cement. Mice recovered for 7–10 days while being habituated to head-fixation.

### CS-US pairing using two distinct tones

Recordings were performed across three experimental sessions: pre-pairing, pairing, and post-pairing lasted between 50-70 minutes. One of CSs consisted of a train of five 3 kHz pure tone pips (70 dB, 50 ms each) and other CS stimulus consisted of five 12 kHz pure tone pips (70 dB, 50 ms each), both delivered at 1 s intervals (pip onsets at 0, 1, 2, 3, and 4 s from trial onset). The identity of CS+ (always co-presented with the shock during pairing) and CS− (3 kHz vs. 12 kHz) was counterbalanced across animals. During pre-pairing and post-pairing, CS+ and CS− were each presented 10 times in an alternating order with a randomized inter-stimulus interval (40–60 s) without any US. During the pairing session, the CS+ co-terminated with a tail shock US (0.7 mA, 10 ms) delivered at 4.5 s (0.5 s after the last pip); the CS− was presented alone. Twenty CS+/US pairings and 20 CS− trials were delivered in the same alternating order (inter-stimulus interval 40–60 s). During post-pairing, CS+ and CS− were presented again without the US in the same format as pre-pairing.

### Behavioral Recordings and Analysis

Whisker and body motion were captured with a high-speed camera positioned to view the head-fixed mouse from the side. For each frame, body and whisker motion were quantified separately from user-defined regions of interest (ROIs) as the sum of absolute pixel-to-pixel differences between consecutive frames (motion energy). Raw motion signals were normalized across the full recording session: (raw − mean) / max(SD, ε).

Trial-by-trial signals were binned at 0.01 s and z-scored relative to a per-trial baseline window of [−10, −1.5] s from CS onset. Three analysis windows were defined: pre-CS [−1.5, 0] s, CS [0, 4.5] s, and post-US/post-CS [4.5, 6] s. Mean z-scores within each window were compared between CS+ and CS− using paired t-tests.

Animals were classified as passive or active defenders based on their behavioral responses during the pairing and post-pairing sessions, assessed from video recordings. Passive defenders (n = 14) displayed suppression of movement and whisker retraction consistent with freezing behavior, without active avoidance or escape attempts. Active defenders (n = 4) showed vigorous wheel running or escape-directed movements during CS+ and US presentations. Classification was performed offline prior to any neural analysis.

### Electrophysiology and Optogenetics

Acute silicon-probe recordings lasted approximately one hour, enabling high-fidelity tracking of most recorded units throughout the experiment. Although this approach precludes longitudinal monitoring of neuronal activity across days, it is well suited for examining sensory stimulus-evoked spiking responses. Silicon probes (P1/H15, 64/128-channel; Cambridge NeuroTech Ltd, Cambridge, UK) were coated with DiI (Thermo Fisher) for visualization of the probe tracks and inserted into the right amygdala of awake, head-fixed mice. Two probe models were used. The P-1 probe carries 64 recording channels distributed across 4 single-sided shanks (16 channels per shank), with electrodes arranged in 2 columns of 8 per shank; row-to-row spacing is 25 µm and column-to-column spacing is 22.5 µm, giving an active recording span of 200 µm per shank. Shanks are spaced 250 µm apart (centre-to-centre), for a total lateral span of approximately 820 µm; each shank is 70 µm wide and 6 mm long. The H15 probe carries 128 recording channels across 2 single-sided shanks (64 channels per shank), with electrodes in 2 columns of 32; spacing is identical (25 µm row-to-row, 22.5 µm column-to-column), giving an 800 µm active span per shank. Shanks are 500 µm apart, 76 µm wide, and 9 mm long. All electrode sites have a manufacturer-specified impedance of approximately 50 kΩ. Signals were acquired at 30 kHz (Intan RHD2000, Intan Technologies, Los Angeles, CA), filtered (0.3–6 kHz), and sorted with Kilosort2.5 (Pachitariu et al., 2024). Clusters were manually curated in Phy (https://github.com/cortex-lab/phy). Clusters with refractory period violations >0.5% were excluded. Only neurons with stable firing rates across the recording sessions were included in the analysis. Units in LA and BA were classified as putative principal neurons (PNs; trough-to-peak > 0.4 ms) or fast-spiking interneurons (INs; trough-to-peak < 0.4 ms, firing rate > 10 Hz).

For sensory stimulations, auditory (3 kHz tone, 70 dB, 50 ms) and tail shock (0.7 mA, 10 ms) stimuli were delivered pseudorandomly (50-100 trials/condition, 5-10 s ITI) using custom MATLAB/Arduino software. For optogenetic inhibition, a 635 nm laser (10 mW/mm^2^, CNI Laser) was delivered through the optic fiber to activate ArchT in PV interneurons, with light-on/off trials interleaved. Light-inhibited neurons (optotagged putative PV interneurons) were identified by comparing firing rate in the recent pre-stimulus window (−0.5 to 0 s) to the earlier baseline (−5 to −0.5 s) using Wilcoxon rank-sum test; neurons were classified as light-inhibited if p < 0.05 and firing rate drop was ≥50%.

Across 24 recording sessions from 23 animals in the tone/shock stimulus presentation experiment (Figures 1–5), automated spike sorting followed by manual curation yielded 938 well-isolated single units (64-channel: mean ± SD: 30 ± 14 per session, range: 12–59, 0.47 units/channel; 128-channel: mean ± SD: 52 ± 25 per session, range: 11–88, 0.41 units/channel). Across 18 recording sessions from 18 animals in the tone-shock pairing experiment (Figures 7–9), automated spike sorting followed by manual curation yielded 1063 well-isolated single units (64-channel: mean ± SD: 49 ± 20 per session, range: 28–87, 0.76 units/channel; 128-channel: mean ± SD: 86 ± 59 per session, range: 26–152, 0.67 units/channel).

**Figure 1.**
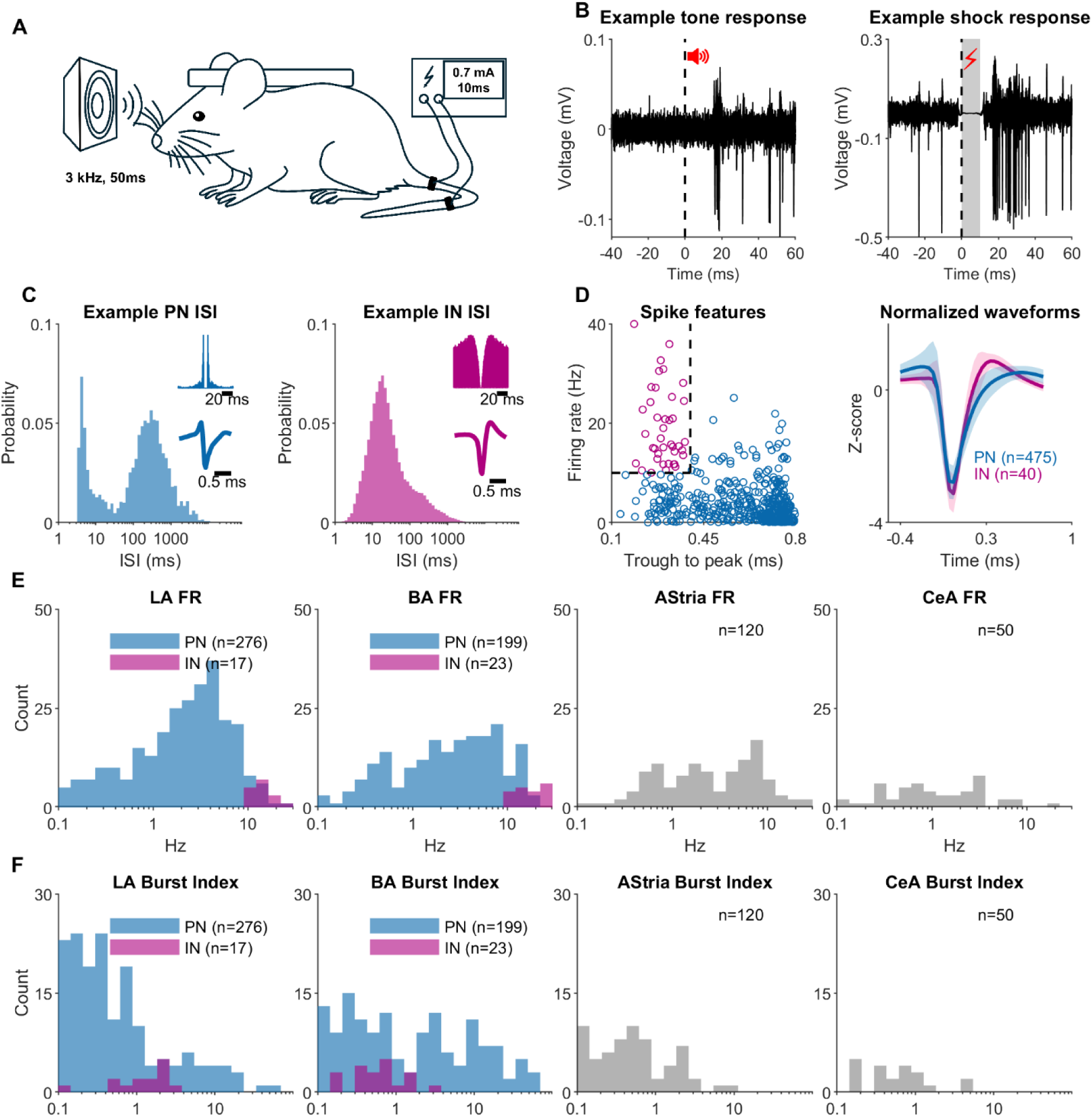
Spike characteristics of neurons in four amygdala regions. **(A)** Experimental setup schematic showing mouse with headplate implantation, auditory stimulus (tone; 3 kHz, 50 ms tone), and aversive stimulus (shock; 10 ms tail-shock). **(B)** Example neuronal responses to tone (left) and shock (right). Top: raw voltage traces showing single-trial responses (5 tone and 6 shock trials superimposed). Red signs indicate the stimulus types, dashed lines the stimulus onset (time 0), and gray shading shows the shock-evoked artefact duration removed before analysis. **(C)** Example interspike interval (ISI) distributions for a putative principal neuron (PN, left, blue) and a fast-spiking interneuron with narrow spike width (IN, right, magenta). Insets show autocorrelograms (top), and average waveforms (bottom). **(D)** Spike feature classification. Left: scatter plot of trough-to-peak time versus firing rate for LA and BA neurons. Dashed lines indicate classification boundaries (0.4 ms and 10 Hz). Right: normalized waveforms (z-score) for PNs (blue, n=475) and INs (magenta, n=40), showing mean ± SD. **(E)** Firing rate distributions across brain regions (LA, BA, AStria, CeA). Plots for LA and BA show separate distributions for PNs (blue) and INs (magenta). Plots for AStria and CeA neurons show results for all neurons (gray). Log-spaced bins from 0.1 to 30 Hz. **(F)** Burst index distributions (Royer et al., 2012) across brain regions. Same color scheme as panel E. Log-spaced bins from 0.1 to 100.

**Figure 2.**
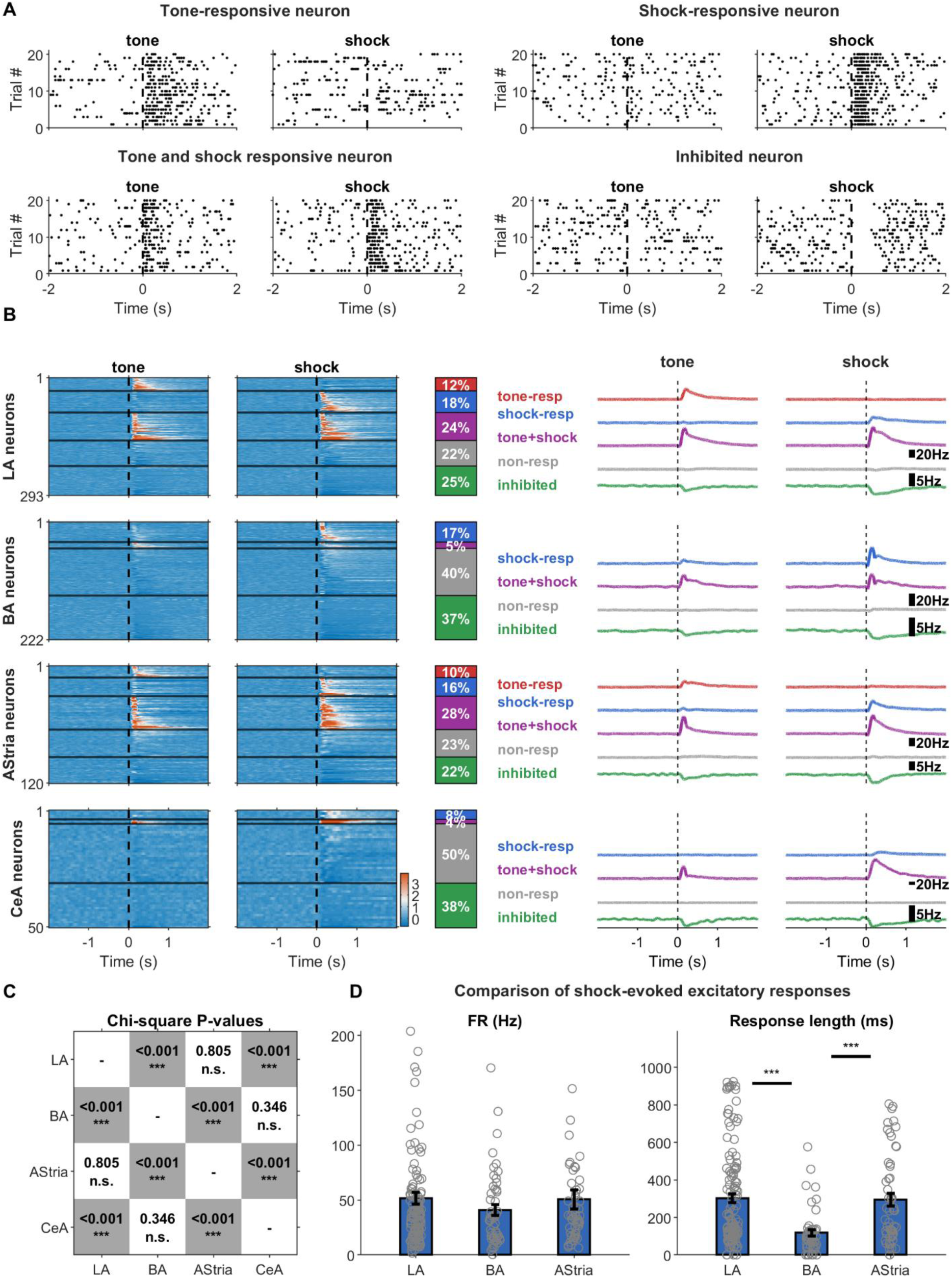
Regional differences in neuronal response profiles to tone and shock stimuli. **(A)** Example single-neuron raster plots for tone (left) and shock (right) stimuli. Top to bottom: tone-responsive neuron, shock-responsive neuron, tone+shock-responsive neuron, and inhibited neuron. Dashed vertical lines mark stimulus onset. **(B)** Population responses across four brain regions (LA, BA, AStria, CeA). Columns 1-2: Z-scored heatmaps for tone and shock trials. Column 3: Bar charts showing cluster proportions (tone-responsive: red, shock-responsive: blue, tone+shock-responsive: purple, non-responsive: gray, inhibited: green). Columns 4-5: Average firing rate (Hz) for tone and shock trials, showing mean response for each cluster with Savitzky-Golay smoothing (201 bins). **(C)** Chi-square test p-value matrix comparing cluster distributions across regions. Gray indicates significant differences. Note that the ratio of cluster proportions is similar between LA and AStria and between BA and CeA. **(D)** Comparison of shock-evoked excitatory response metrics across LA, BA, and AStria. Bar graphs show no difference in firing rate (FR) but a difference in response length for shock-responsive and tone+shock-responsive neurons combined. Individual data points shown as circles; bars represent mean ± SEM. For results see Table S3.

**Figure 3.**
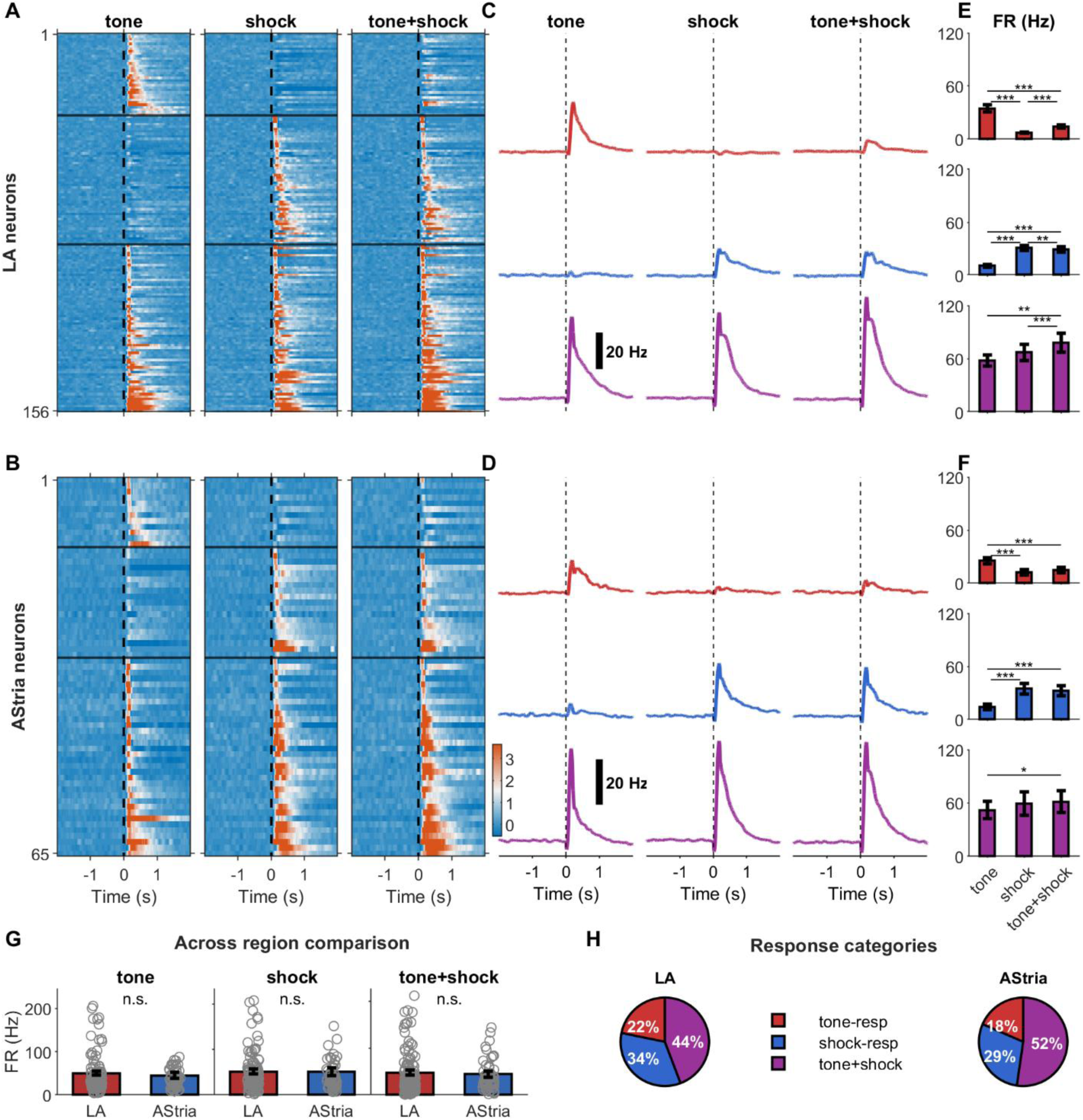
Similar firing rate in tone-responsive, shock-responsive, and tone+shock-responsive neurons in the LA and AStria. **(A)** Heatmaps for LA neurons showing z-scored peristimulus time histograms for tone (left), shock (middle), and tone+shock (right) stimuli. Neurons sorted by cluster (tone-responsive, shock-responsive, tone+shock-responsive) based on tone and shock responses only. Color scale shows z-score normalized to baseline. Black lines separate clusters. **(B)** Heatmaps for AStria neurons, same format as panel A. **(C)** Lines showing population average firing rates (Hz) for tone-responsive (red), shock-responsive (blue), and tone+shock-responsive (purple) neurons in the LA. Each cluster displayed separately (stacked rows) for tone, shock, and tone+shock stimuli. **(D)** Lines for AStria neurons, same format as panel C. **(E)** Bar graphs showing firing rate (FR) comparing tone, shock, and tone + shock responses in LA neurons within each cluster. Bars show mean ± SEM. For results see Table S4. **(F)** Bar charts for AStria neurons, same format as panel E. For results see Table S4. **(G)** Firing rate (FR) for tone (left), shock (middle), and tone+shock (right) responsive neurons in the LA and AStria is similar. Individual data points shown as gray circles. Y-axis scaled to 95th percentile to avoid compression from outliers. For results see Table S5. **(H)** Similar proportions of tone-responsive, shock-responsive, and tone+shock-responsive neurons in LA (left) and Astria (right). Percentages indicate fraction of responsive neurons in each category.

**Figure 4.**
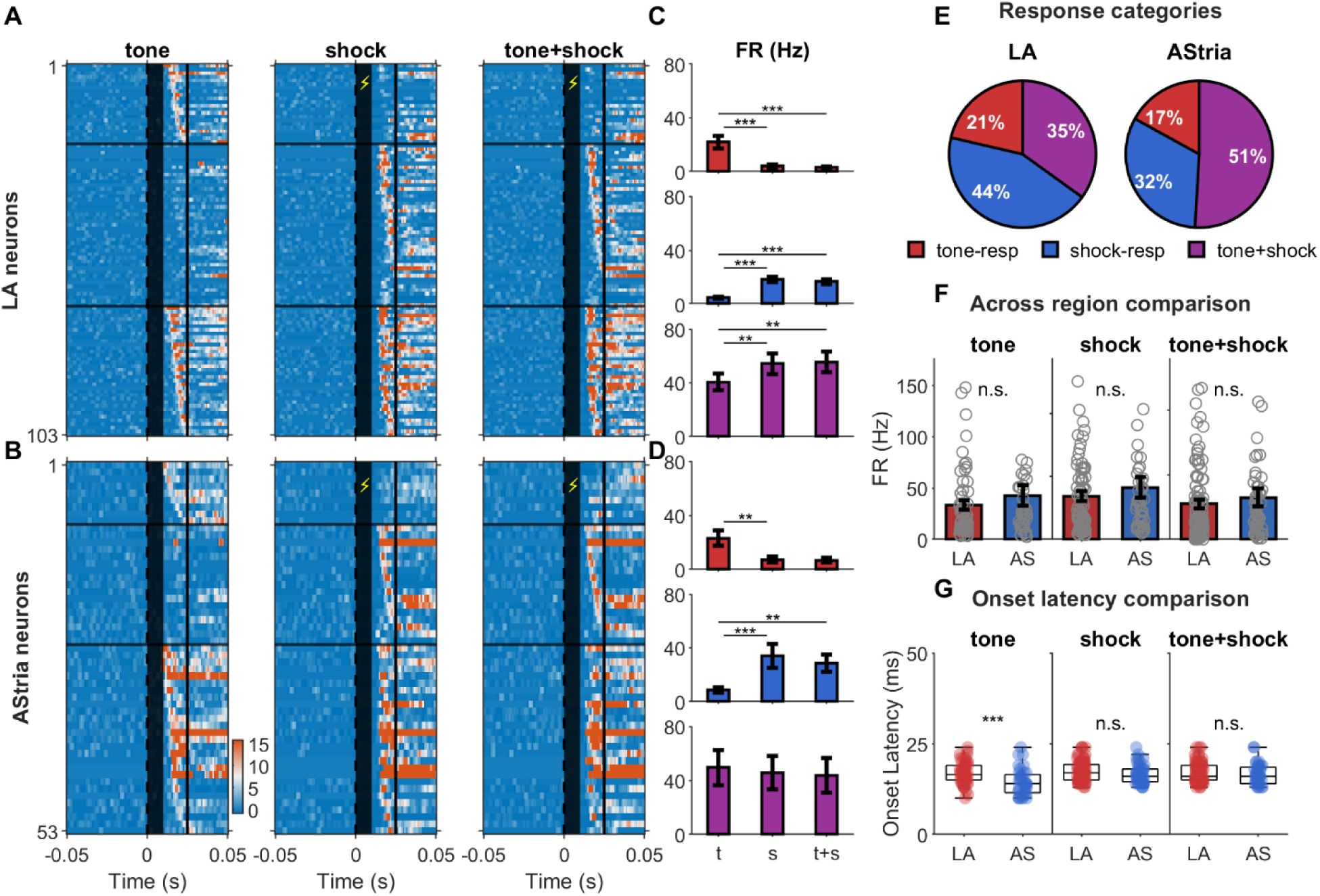
Monosynaptic responses to tone and shock stimuli are similar in the LA and AStria. **(A)** Heatmaps of monosynaptic responses in LA neurons showing z-scored peristimulus time histograms for tone (left), shock (middle), and tone+shock (right) stimuli. Only neurons with detected responses in the monosynaptic window (0-25ms post-stimulus) are shown, sorted by cluster (tone-responsive, shock-responsive, tone+shock-responsive). Color scale shows z-score normalized to baseline. Black lines separate clusters. **(B)** Heatmaps for AStria neurons, same format as panel A. **(C)** Firing rate (FR) bar charts comparing tone, shock, and tone+shock responses for each LA cluster. Stacked vertically: tone-responsive (top), shock-responsive (middle), tone+shock-responsive (bottom). Bars show mean ± SEM. For results see Table S7. **(D)** Firing rate (FR) bar charts for AStria neurons, same format as panel C. **(E)** Pie charts showing similar proportions of tone-responsive, shock-responsive, and tone+shock-responsive neurons among monosynaptic responders in the LA (left) and AStria (right). Percentages indicate fraction of monosynaptic neurons in each category. **(F)** Across region comparison of firing rate (FR) for tone (left), shock (middle), and tone+shock (right) monosynaptic responders comparing LA vs AStria neurons. Individual data points are shown as gray circles. Bars show mean ± SEM. Y-axis scaled to 95th percentile to avoid compression from outliers. For results see Table S8. **(G)** Onset latency comparison for monosynaptic responses. Three plots show tone, shock, and tone+shock onset latencies comparing LA vs AStria neurons. Bars show mean ± SEM. For results see Table S8. Statistical comparison using Wilcoxon rank-sum test (p<0.05, p<0.01, **p<0.001, n.s.=not significant).

**Figure 5.**
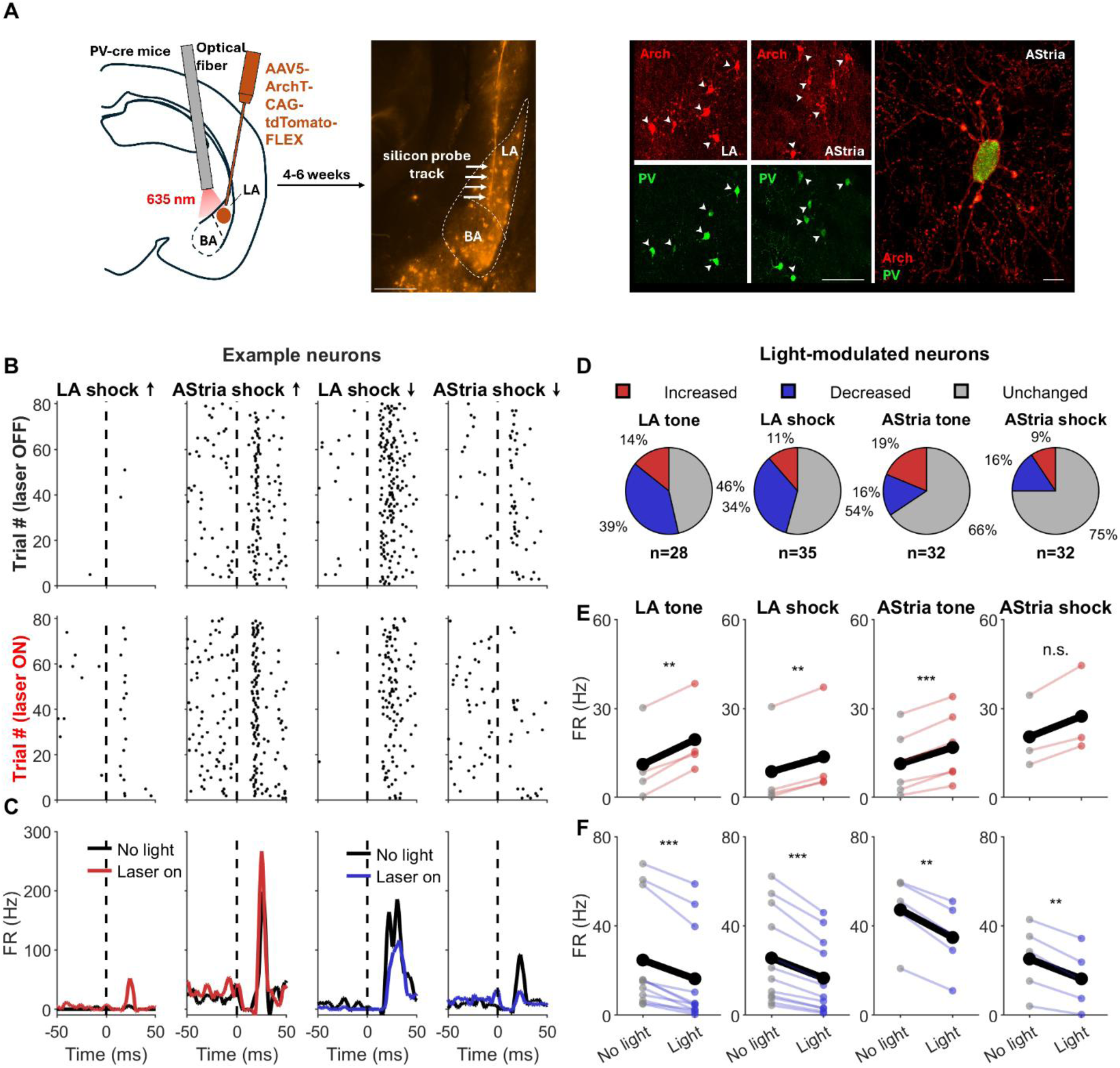
A fraction of monosynaptic sensory stimulus-evoked spike responses in the LA and AStria are controlled by PV interneurons. **(A)** Left: Schematic of stereotaxic injections in PV-Cre mice. AAV5-CAG-FLEX-ArchT-tdTomato virus was delivered into the LA and AStria, followed by implantation of an optical fiber positioned above the LA or AStria for 635-nm light delivery to inhibit PV-expressing interneurons. Following 4–6 weeks of viral expression, a silicon probe was inserted to record spiking activity during optogenetic manipulation. Right: Representative fluorescence image showing tdTomato-expressing neurons in the BA and LA, along with the track of the silicon probe (arrows). Left: Micrographs showing tdTomato-expression (top) in PV interneurons (bottom) in the LA and AStria. Right: A PV interneuron in Astria expresses ArchT throughout the membrane indicated by tdTomato signal. Scale bars: left, 100 μm; right, 10 μm. **(B)** Example raster plots showing individual neuron responses to tone or shock stimuli without light (top row) and with light (middle row) for four example neurons. Each row represents one trial, with spike times shown as dots. Black dashed vertical line indicates stimulus onset. **(C)** Lines for the same example neurons showing average firing rate (FR) over time for no-light (black) and light (color) conditions. Time axis centered at stimulus onset (0 ms). Each plot shows the peristimulus time histogram comparing baseline and optogenetic manipulation conditions. **(D)** Pie charts showing proportions of neurons with enhanced, decreased, and unchanged monosynaptic responses during inhibiting PV interneurons. Numbers and percentages indicate fraction of responsive neurons in each category. **(E)** Plots showing trial-averaged spike counts for individual neurons with increased spiking comparing no-light (left, black) vs light (right, red) conditions. Each line represents one neuron, with population mean shown in bold. Four panels correspond to LA tone, LA shock, AStria tone, and AStria shock enhanced neurons. Enhancement detected using Wilcoxon signed-rank test (p<0.05) comparing spike counts in the monosynaptic window (12-50 ms) between light and no-light conditions. For results see Table S9. **(F)** Plots showing trial-averaged spike counts for individual neurons with decreased spiking comparing no-light vs light conditions, formatted as in panel F. Decreased responses were identified using the same statistical criteria. For results see Table S9.

### Data Analysis and Statistics

Analyses were performed with custom MATLAB scripts (R2024b). Peristimulus time histograms (1 ms bins, −5 to +4 s from stimulus) were smoothed with a Savitzky-Golay filter (201 ms for population, 5 ms for monosynaptic analysis) and delay-corrected. Baselines were from −5 to 0 s. The burst index (adapted from (Royer et al., 2012) was calculated using the CellExplorer framework (Petersen et al., 2021). To quantify bursting propensity, we computed the ratio of the mean autocorrelogram value (3–5 ms) to the baseline (mean of 200–300 ms). Values > 1 indicate a higher probability of short-interval spikes (bursts) relative to tonic firing.

#### Neuronal Classification

Neurons were classified based on z-scored responses. For population analysis (Figures 2, 3), excitation was z ≥ 1.5 (0-1 s window) and inhibition was a ≥50% firing rate drop. For monosynaptic analysis (Figure 4), z ≥ 5 and probability ≥ 0.1 was used. The initial 10 ms window after US presentation was excluded. Functionally, neurons were clustered as tone-responsive, shock-responsive, tone+shock-responsive, non-responsive, or inhibited.

#### CS+/CS− Plasticity Classification

For neurons recorded across all three experimental sessions related to the CS-US pairing, plasticity was quantified as the change in mean z-scored response from pre-pairing to post-pairing. Z-scores were computed relative to a [−8, 0] s baseline window using 5 ms bins, with Savitzky-Golay smoothing (101 bins) and delay correction applied before z-scoring. For each neuron, ΔCS+ was defined as the mean z-score in the [0, 5] s post-stimulus window during the post-pairing session minus the corresponding value from the pre-pairing session; ΔCS− was computed identically for CS− trials. Neurons were classified into five classes based on the direction and stimulus-specificity of the change: CS+ up (CS+↑: ΔCS+ > 0.2, exceeding ΔCS− by >0.1), CS+ down (CS+↓: ΔCS+ < −0.2, more negative than ΔCS− by >0.1), CS−up and CS− down (CS−↑ and CS−↓: same logic applied to ΔCS−), and Stable (all remaining neurons). A subset of CS+ down neurons whose post-pairing CS+ response fell below baseline (z < −0.1) were designated CS+ down inhibited and treated as a separate subclass in group-level analyses (Figure 8). Neurons with insufficient spiking activity (<5 spikes in <5 trials across any recording phase) were assigned to the Stable class.

#### Trial-by-Trial Response Analysis

Per-trial z-scores were computed for each neuron across all three experimental sessions by binning spikes at 5 ms, smoothing with a Savitzky-Golay filter (101 bins), applying delay correction, converting to firing rate (Hz), then z-scoring against the [−8, 0] s baseline. Mean z-scores were extracted within the following windows: CS response [0, 5] s for pre-pairing and post-pairing; CS response [0, 4.5] s and US response [4.5, 5] s for the pairing session. Within-session CS+ versus CS− comparisons were performed using Wilcoxon signed-rank test.

#### US Responsiveness Detection

Within the CS+ up class (class 1), individual neurons were further classified as US-responsive or non-US-responsive based on their firing during the first 2 pairing trials. A neuron was classified as US-responsive if it showed a significant increase in firing (z ≥ 1.5 above CS-level activity, sustained for ≥0.5 s) within 2 s of US onset. Neurons not meeting these criteria were classified as non-US-responsive, and their US-evoked activity was quantified from a fixed [4.5, 5.5] s window. Early-versus-late pairing comparisons (first 2 vs. last 2 trials) were performed using Wilcoxon signed-rank test.

#### Firing features during pre-pairing and US responsiveness

Baseline firing rate was estimated from the pre-pairing session as the mean spike count in the [−8, 0] s window divided by trial count and window duration. US responsiveness was the mean per-trial z-score in the [4.5, 5.5] s window during CS-US pairing. Burst index values were obtained from the CellExplorer framework (burstIndex from (Royer et al., 2012). Differences between plasticity groups were assessed with Kruskal-Wallis ANOVA; pairwise Wilcoxon rank-sum test with Bonferroni correction were performed when Kruskal-Wallis ANOVA revealed p < 0.05.

#### Statistical Comparisons

Proportions were compared using Chi-square test with Cramér’s V effect size. Non-normal distributions (firing rates, latencies, durations) were compared using Wilcoxon rank-sum test (for two groups) or Kruskal-Wallis ANOVA (for three or more groups) followed by Dunn’s post-hoc test. Repeated measures (e.g., tone vs. shock vs. tone+shock responses) were analyzed using Friedman ANOVA with post-hoc Wilcoxon signed-rank test. For optogenetics, light-on/off trials were compared using a Wilcoxon signed-rank test or paired t-test. Data are reported as mean ± SEM, except for baseline spike feature characterizations which are mean ± SD. Statistical significance was defined as p < 0.05.

### Immunohistochemistry and imaging

To verify viral expression and electrode placement, mice were anesthetized with isoflurane after the recordings and decapitated. Brains were removed, post-fixed for 24 hours in 4% paraformaldehyde, and sectioned coronally (50 µm) on a Leica VT1000S vibratome. Immunostaining was used to verify the specificity of ArchT expression in PV interneurons. Sections were blocked using 10% Normal Donkey Serum in 0.5% Triton X-100 dissolved in 0.1 M PB and incubated for 4 days at 4°C with primary antibody (guinea pig anti-PV, 1:10,000; Synaptic Systems, 195 004). Sections were then incubated with secondary antibody (donkey anti-guinea pig Alexa 488, 1:1000; Jackson ImmunoResearch, 706-545-148) overnight at room temperature. Images were acquired using a confocal microscope (Leica Stellaris 8 Falcon Scanning Confocal) to confirm overlap between ArchT-tdTomato expression and PV immunolabeling.

### Histology and code accessibility

Probe placement was confirmed histologically on 150 µm coronal sections based on the Dil signal. Only data from on-target animals were included. When a probe shank spanned more than one amygdala sub-area, units were assigned to sub-regions based on the identity of the channel that yielded the highest-amplitude spike waveform for each unit (the best channel, determined during manual curation in Phy). Because electrode positions along the shank are fixed and precisely known from the probe geometry (25 µm centre-to-centre spacing), each best channel maps to an unambiguous dorso-ventral coordinate. Sub-regional boundaries were identified independently from DiI-labeled coronal sections aligned to mouse brain reference atlas of Allen Institute for Brain Science (https://mouse.brain-map.org/static/atlas), and the corresponding boundary depth was back-projected onto the electrode map. Units whose best channel fell within 25 µm of a confirmed sub-regional boundary were excluded from region-specific analyses. Analysis scripts are available from the corresponding author upon reasonable request.

## Results

### Baseline firing features of neurons in four amygdala regions

To reveal the spike responses in amygdala circuits during stimulus processing, we recorded single-unit activity in 23 mice across four subregions (LA, n = 293; BA, n = 222; AStria, n = 120 and CeA, n = 50) in awake, head-fixed mice (Figure 1A). Electrode placement was reconstructed post hoc by visualizing the silicon probe tracks using DiI, confirming the recording sites being in the LA, BA, AStria, and CeA (Figure S1, B). In the LA and BA, units were classified as putative principal neurons (PNs) or putative fast-spiking interneurons (INs) based on spike waveform features and autocorrelograms (Figure 1C, D). PNs exhibited significantly longer spike trough-to-peak durations and lower firing rates compared to INs (Table S1). In the CeA and AStria, local inhibitory cells could not be separated from medium spiny neurons based on these two spiking features, therefore, all units in these two regions were pooled together. Overall, our baseline characterization of firing across these four subregions revealed that LA and BA INs fired with the highest rate while CeA neurons fired with the lowest rate, out of all studied cell types. In addition, BA PNs had a higher burst index than LA PNs, AStria and CeA neurons, while LA and BA INs showed higher burstiness than AStria and CeA neurons (Figure 1E, F; Table S1). Lastly, we inspected the response stability during the experiments by analyzing the changes in firing rates across blocks and confirmed consistent activity levels throughout the recording sessions (Figure S2; Table S2).

### Similar firing dynamics in LA and AStria characterize auditory and aversive stimulus-evoked responses

The canonical model of the amygdala states that the LA and BA function as a tightly coupled unit, where sensory information integrated in the LA is relayed to the BA before transmission to output nuclei like the CeA (Johansen et al., 2011; Tovote et al., 2015; Trent et al., 2025). We tested this hypothesis by comparing subregional responses to the tone (auditory stimulus) and shock (aversive stimulus) (Figure 1B). We classified neurons into distinct functional types: tone-responsive, shock-responsive, tone+shock-responsive, non-responsive, or inhibited (Figure 2A, B). In the LA, we observed a diverse distribution of responsiveness: tone-responsive (12%, n = 34), shock-responsive (18%, n = 53), tone+shock-responsive (24%, n = 69), non-responsive (22%, n = 63), and inhibited (25%, n = 74). This profile was mirrored in the AStria, which showed a similar composition: tone-responsive (10%, n = 12), shock-responsive (16%, n = 19), tone+shock-responsive (28%, n = 34), non-responsive (23%, n = 28), and inhibited (23%, n = 27). In contrast, the BA was dominated by non-responsive (40%, n = 88) and inhibited (38%, n = 83) neurons, with a shock-responsive population (17%, n = 37) resembling the LA and AStria and a minor group of tone+shock-responsive units (5%, n = 11). Similarly, the CeA was largely quiescent, primarily comprised of non-responsive (50%, n = 25) and inhibited (38%, n = 19) cells, with smaller populations of shock-responsive (8%, n = 4) and tone+shock-responsive (4%, n = 2) neurons. Our data strengthen the dissociation between the LA and BA in sensory stimulus processing, while highlighting that the LA and AStria may similarly process sensory stimulations.

To corroborate this conclusion, we compared the proportions of the five response categories among the four regions. We found that the LA and AStria exhibited statistically indistinguishable distributions (Chi-square test, p = 0.805, Cramér’s V = 0.06; Figure 2C). In contrast, both the BA and CeA showed significantly lower responsiveness that was comparable in these two nuclei (Chi-square test, p = 0.371, Cramér’s V = 0.14; Figure 2C) but was different from the LA and AStria (Chi-square test, p < 0.001, Cramér’s V = 0.23; Figure 2C). As shock evoked spiking in a similar ratio of neurons in the LA, BA and AStria, we wondered whether the response properties of shock-excited neurons were similar in these regions. We found that the rate of shock-driven firing was robust and matched across LA, AStria, and BA, while responses in BA neurons were significantly shorter in duration (Figure 2D, Table S3). These findings further suggest that, under these conditions, the LA and AStria - not the LA and BA - operate similarly as circuits for processing sensory stimuli.

### LA and AStria process sensory information in parallel

Next, we tested how the LA and AStria integrate sensory information by analyzing excitatory responses to simultaneous tone and shock presentation in the responsive neuronal clusters. We observed that tone-responsive neurons exhibited significant suppression both in firing rate and response length during combined stimulation in both regions (Figure 3A-F, Table S4). In contrast, shock-responsive neurons maintained robust firing during co-presentation of tone+shock, with AStria responses showing no significant change in firing rate or response length and LA responses showing only a slight reduction (Figure 3A-F, Table S4). In tone+shock-responsive neurons, we found evidence of supralinear summation analyzed in this long (<500 ms) time window. In the LA, the neuronal response to the combined stimuli significantly exceeded individual tone- and shock-driven responses (both firing rate and response length), while, in the AStria, the tone+shock co-presentation resulted in a spiking that was elevated only compared to tone-driven firing (Figure 3A-F, Table S4). If we restricted the analysis to LA PNs, the results recapitulated the population data obtained by analyzing the spike responses in the three groups (Figure S3A, B). In contrast, LA INs, which accounted for only 8% of all sensory stimulus-excited LA neurons, were found exclusively among neurons responsive to shock or tone+shock stimulation (Figure S3C-E). Crucially, a direct comparison of the firing rate and response length across all responsive neurons confirmed that signal gain was comparable across the LA and AStria for tone, shock, and combined stimuli (Figure 3G, Table S5). The spike elevations were observed in similar proportions of responsive cell types (Chi-square test, χ^2^ = 1.20, p = 0.555, Cramér’s V = 0.07; Figure 3H), supporting the hypothesis of shared integration properties.

### Sensory stimuli preferentially excite fast spiking inhibitory cells in BA

In the BA, only a relatively small fraction of neurons was excited by sensory stimuli, whereas a large proportion was inhibited (Figure 2B). This observation raised the possibility that sensory stimuli preferentially recruit local INs. To test this hypothesis, we compared stimulus-evoked spiking in BA PNs and BA INs. We found that BA INs accounted for 21.3% of all sensory stimulus-excited BA neurons (10 of 47 neurons, Figure S4A, B, G). Furthermore, BA INs exhibited significantly larger increases in firing in response to shock and tone+shock stimulations than BA PNs (Figure S4F, Table S6). Notably, excitatory responses in both PNs and INs were most commonly evoked by the shock alone or by the combined tone+shock stimulus, whereas tone-only responses were rare. These findings suggest that tone-evoked activity originating in the LA is generally insufficient to drive spiking in downstream BA neurons.

To further examine the recruitment of fast-spiking inhibitory neurons, we identified parvalbumin interneurons (PV INs) using optotagging (Figure S4C). A Cre-dependent inhibitory opsin (AAV5-CAG-FLEX-ArchT-tdTomato) was injected into the BLA of PV-Cre mice (n = 4), and silicon probe recordings were performed in the BA. Among stimulus-excited BA neurons (n = 13), seven were identified as optotagged PV INs (54%, Figure S4D, E, G). Stimulus-evoked firing increases were generally larger in PV INs than in simultaneously recorded other BA neurons, although the difference was statistically significant only during tone+shock co-stimulation (Figure S4F, Table S6).

Given that PV INs constitute only approximately 6% of the total BA neuronal population (Vereczki et al., 2021), their disproportionately high representation among stimulus-excited neurons suggests that these inhibitory cells are preferentially recruited by sensory stimuli within BA circuits. Although the number of recorded neurons was insufficient to directly determine whether PV INs mediate the widespread inhibition observed in the BA population (Figure 2), their preferential recruitment and robust stimulus-evoked firing support the idea that they contribute to suppressing spiking activity in BA neurons.

### Parallel monosynaptic drive to the LA and AStria networks

The similar excitatory activity in LA and AStria neurons driven by tone and shock raises the question of their afferent connectivity. As previous studies showed that the inputs from the thalamus innervate both the LA and AStria (LeDoux et al., 1985; LeDoux et al., 1991; Barsy et al., 2020), we analyzed the onset latencies of spikes to identify monosynaptic responses (<25 ms) indicative of direct and parallel thalamic drive (Figure 4A, B)(Quirk et al., 1995; Quirk et al., 1997). During the analysis we found a significant population of neurons that emitted short-latency spikes in both regions, and similarly to large time scale dynamics, monosynaptic integration varied by cell type. While tone-responsive neurons exhibited profound suppression during combined stimulation (Figure 4C, D, Table S7), shock-responsive neurons maintained elevated firing (Figure 4C, D, Table S7). In tone+shock-responsive neurons, the combined monosynaptic response and shock-driven monosynaptic responses were similar, showing no supralinear summation (Figure 4C, D, Table S7). Comparable proportions of responsive cell types were observed in the LA and AStria (Chi-square test, χ^2^ = 3.74, p = 0.159, Cramér’s V = 0.15, Figure 4E), supporting the hypothesis of shared integration properties. In addition, the firing rate of these monosynaptic responses was comparable across stimuli in both nuclei (Figure 4F, Table S8). These data show that monosynaptic excitation evokes a similar increase in firing in LA and AStria neurons as we observed at longer timescales (Figure 3). The only difference noticed was the absence of supralinear summation, indicating that the building network activity upon tone and shock co-presentation should be responsible for supralinear response.

Lastly, we examined the onset latencies of sensory stimulus-evoked spiking. Whereas the onset latencies for shock responses were statistically indistinguishable between regions, tone responses were significantly faster in the AStria compared to the LA (Figure 4G, Table S8). This spike timing, with AStria responding prior to or concurrently with the LA, rules out a simple serial relay from LA to AStria at short-latency firing. Further, the results showing shorter onset latencies but similar firing rates in AStria compared to LA also imply that tone and shock information should be conveyed via distinct afferents at least to the AStria. This conclusion is further supported by the fact that tone-evoked response latencies in AStria neurons were shorter than shock-evoked response latencies (tone: 14.4 ± 0.6 ms, n = 36; shock: 16.6 ± 0.4 ms, n = 44; Wilcoxon rank-sum test, p = 0.001).

### PV interneurons control spiking in the LA and Astria in a feedforward manner

As thalamic inputs drive feedforward excitation and inhibition in parallel, we next asked whether this functional similarity also extends to local inhibitory regulation in both LA and AStria. PV INs providing powerful feedforward inhibition in the LA are considered the primary interneurons regulating network and subnetwork activities (Wolff et al., 2014; Lucas et al., 2016). To determine if PV IN-mediated feedforward inhibition is conserved in the AStria, we targeted PV INs in PV-Cre mice (n = 12) by injecting AAV5-CAG-FLEX-ArchT-tdTomato into the amygdala region (Figure 5A). Immunocytochemical analysis confirmed specific expression of ArchT in PV INs in both the LA and AStria (Figure 5A). To reveal the function of PV INs in controlling short-latency spiking, we optogenetically silenced these inhibitory cells during stimulus presentation. First, we confirmed the efficacy of PV IN inhibition. Here, we compared their firing rates during the pre-stimulus period (−0.5 s to 0 s) with and without light delivery (Figure S5A). In light-inhibited neurons, i.e. in PV Ins, laser illumination significantly suppressed firing rates in both regions during this 0.5 s-long time window (LA: 4.59 ± 0.91 Hz no-light vs 1.69 ± 0.38 Hz light, n=16, p < 0.001; AStria: 5.69 ± 0.93 Hz no-light vs 2.02 ± 0.33 Hz light, n=6, p < 0.001; Wilcoxon signed-rank test; Figure S5B-E). This light-induced suppression in PV IN spiking corresponded to a reduction in their mean firing rate of 63.20% in LA and 64.50% in AStria PV. Then, we assessed the inhibitory effect of PV INs on tone- and shock-triggered spiking. We found that inhibiting PV INs resulted in modulated sensory stimulus-triggered responses at the level of single neurons both in the LA and AStria (Figure 5B, C). We observed distinct subpopulations that either increased or decreased their firing rates. The proportion of neurons exhibiting increased, decreased, or unchanged responses was similar in LA and AStria for both tone (Chi-square test, χ^2^ = 4.28, p = 0.117) and shock stimuli (Chi-square test, χ^2^ = 3.48, p = 0.176; Figure 5D). In the LA, a substantial proportion of neurons decreased their response (tone: 39%, n=11; shock: 34%, n=12), exhibiting significant reductions in firing rate (Figure 5E, F, Table S9). A smaller subset showed increased activity (tone: 14%, n=4; shock: 11%, n=4), with significantly elevated firing rates (Figure 5E, F, Table S9). In the AStria, response modulation was also bidirectional, with neurons exhibiting decreased activity (tone: 16%, n=5; shock: 16%, n=5) or increased activity (tone: 19%, n=6; shock: 9%, n=3) (Figure 5E, F, Table S9). The fact that both regions operate with a similar feedforward inhibition further strengthens the hypothesis that the LA and AStria work in parallel during sensory signal processing. Together, these data suggest that the LA and AStria function as a parallel processing unit, sharing input sources, integration properties, and a similar feedforward inhibitory motif.

### CS-US pairing in awake, head-fixed mice

In the next set of experiments, we aimed to determine whether CS–US pairing, comparable to discriminative Pavlovian fear conditioning, induces similar or distinct changes in spiking activity in LA and AStria neurons. Head-fixed mice were exposed to two distinct tones (3 kHz, 12 kHz), each consisting of five beeps. During the pre-pairing (habituation) and post-pairing (recall) sessions, the two tones were presented alternately 10 times each. During the CS–US pairing session, CS+ and CS− were alternately presented 20 times, and each CS+ co-terminated with a US (Figure 6A).

**Figure 6.**
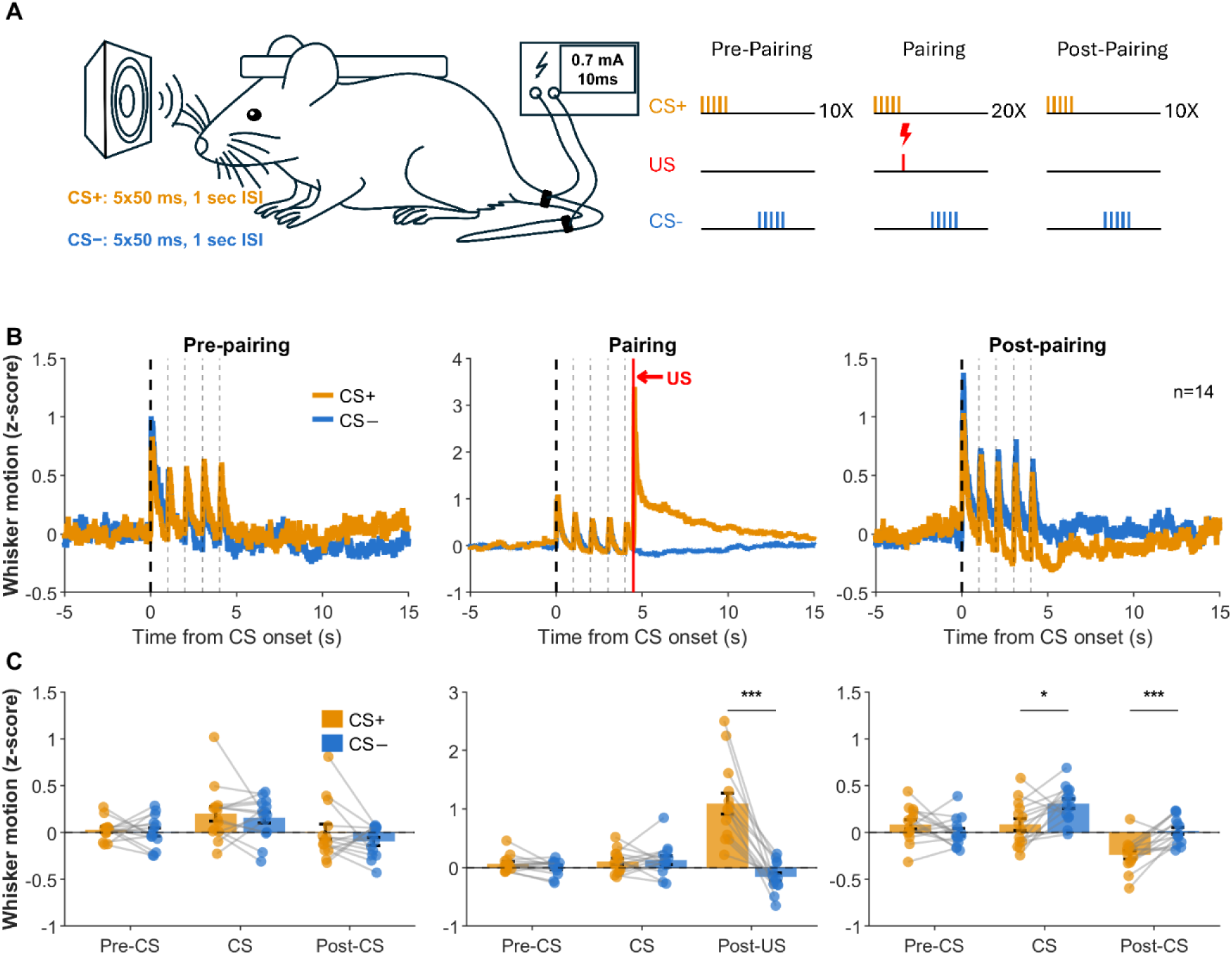
Experimental paradigm and sensory stimulus-evoked whisker motion in passive defenders. **(A)** Experimental design. Left: schematic of the head-fixed recording setup with silicon probe implantation. Right: behavioral paradigm showing the three experimental sessions (pre-pairing, pairing, and post-pairing) and the timing of CS (tone) and US (shock) presentations. **(B)** Mean z-scores of whisker motion for passive defenders (n = 14 animals) across three sessions (pre-pairing, pairing, and post-pairing). CS+ (orange) and CS− (blue) traces are shown separately. Time axis is aligned to CS onset (t = 0 s). **(C)** Bar graphs showing mean ± SEM whisker motion z-scores within pre-CS, CS, and post-CS windows for each session. Individual animal means are overlaid as dots. Paired t-test was used to compare CS+ versus CS− responses within each window and session. For results see Table S10.

By monitoring whisker and body movements throughout the experiment, we identified two distinct behavioral response patterns. Most mice (n = 14) exhibited increased whisking and body movements during US presentations and reduced whisking during both CS+ and post-CS+ periods compared to CS− following CS–US pairing (Figure 6B, C and Figure S6A, Table S10). Body movements were also significantly suppressed during the post-CS+ period (Figure S6A, Table S10). In contrast, a smaller group of mice (n = 4) showed significantly increased whisking and body movements during CS–US pairing, specifically during CS+ and US presentations, as well as during the post-CS+ period compared to post-CS− presentations (Figure S6B, C, Table S10). Based on these behavioral responses, mice were classified as passive defenders, which reduced whisking and body movements in response to CS+, or active defenders, which increased these behaviors.

In the LA, we recorded a sufficient number of units to assess whether neuronal responses to CS–US pairing differed between passive and active defenders. We compared both the magnitude of CS+- and CS−-evoked spiking and the changes in CS+-versus CS−-evoked activity between the post-pairing and pre-pairing sessions. Neither analysis revealed significant differences between passive and active defenders (Figure S7, Table S11). Therefore, recordings from all 18 mice were pooled for subsequent analyses (see below). These findings are consistent with previous studies showing that neuronal representations in the LA encode CS–US associations (Reijmers et al., 2007; Rashid et al., 2016; Abdou et al., 2018; Abatis et al., 2024) but do not necessarily predict the specific behavioral output expressed during defensive responses (Goosens et al., 2003).

### Similar associative plasticity in CS-evoked spiking in the LA and AStria following CS-US pairing

To compare spike responses in LA and AStria neurons associated with CS–US pairing, we recorded spiking during three experimental sessions: pre-pairing (habituation), CS–US pairing, and post-pairing (recall) (Figures 6 and 7). Based on changes in post-pairing responses relative to pre-pairing activity, neurons in both regions could be classified into five categories: neurons that significantly i) increased (CS+ up) or ii) decreased (CS+ down) their firing in response to CS+; neurons that significantly iii) increased (CS− up) or iv) decreased (CS− down) their firing in response to CS−; and v) neurons that showed no significant change in spiking activity (stable) (Figure 7A–D).

Comparison of the magnitude of pairing-induced changes revealed no significant differences between LA and AStria neurons, except for responses of CS+ down neurons during CS− presentations (Figure 7E, Table S12, Figure S8, Table S13). Likewise, the proportions of neurons belonging to the five response categories were similar in the two regions (χ^2^ = 10.28, p = 0.036, Cramér’s V = 0.18, χ^2^ test; Figure 7F, Table S14). These findings indicate that CS–US pairing induces comparable associative plasticity in CS-evoked spiking activity in both the LA and AStria.

**Figure 7.**
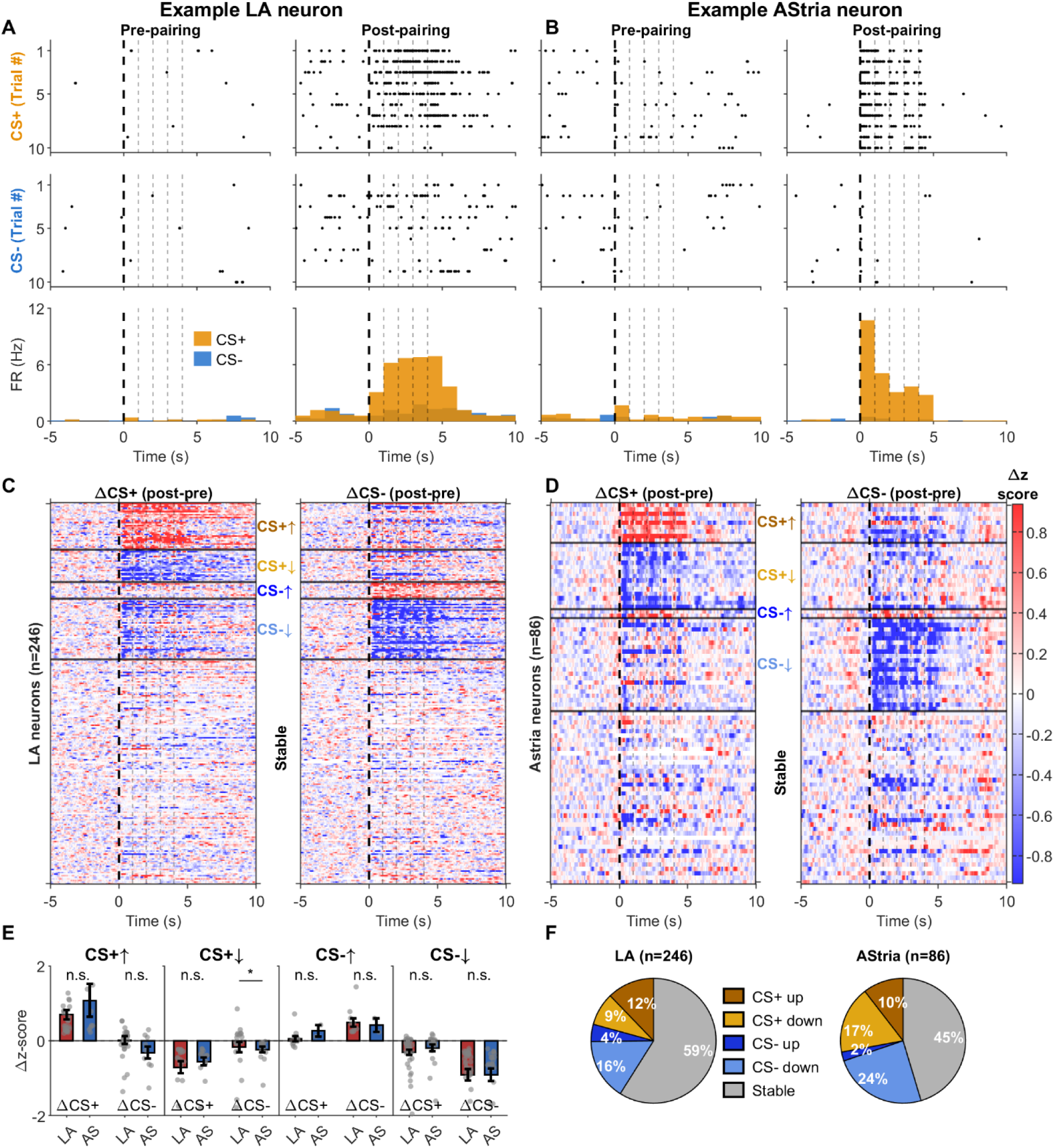
Bidirectional plasticity of CS+ and CS− responses induced by CS-US pairing. **(A, B)** Raster plots and mean firing rate histograms (1000 ms bins) showing CS+ and CS− evoked spiking activity during pre-pairing and post-pairing sessions for representative CS+ up (↑) neurons in LA (A) and AStria (B). CS+ trials are shown in orange; CS− trials in blue. Dashed vertical lines indicate beep onset times (t = 0, 1, 2, 3, 4 s). **(C, D)** Population delta (Δ)-response heatmaps (post-pairing minus pre-pairing, z-score) for CS+ (ΔCS+) and CS− (ΔCS−) responses in LA (C) and AStria (D). Five categories are shown: CS+↑ (CS+ up: increased CS+ response), CS+↓ (CS+ down: decreased CS+ response), CS−↑, CS−↓, and Stable. **(E)** Bar graphs comparing ΔCS+ and ΔCS− between LA and AStria for each plasticity class. Bars show mean ± SEM; individual neuron values are overlaid. Statistical comparison using Wilcoxon rank-sum test. For results see Table S12. **(F)** Pie charts showing the distribution of plasticity classes in LA and AStria. Chi-square test was used to compare class distributions between regions. For results see Table S14.

### Similar dynamics of CS-evoked spiking in the LA and AStria associated with CS-US pairing

Next, we compared the temporal dynamics of spiking activity in CS+ up neurons from the LA and AStria. During the pre-pairing session, these neurons exhibited substantial and highly similar habituation of responses to both CS+ and CS− presentations (Figure 8A, Table S15). During pairing, the initially weak CS-evoked firing gradually recovered after the first few US presentations. This increase was maintained during subsequent CS+ presentations but not during CS− presentations, resulting in a significant divergence between CS+ and CS− responses in both regions. This difference persisted throughout the post-pairing session, indicating the development of discriminative plasticity (Figure 8A, Table S15).

**Figure 8.**
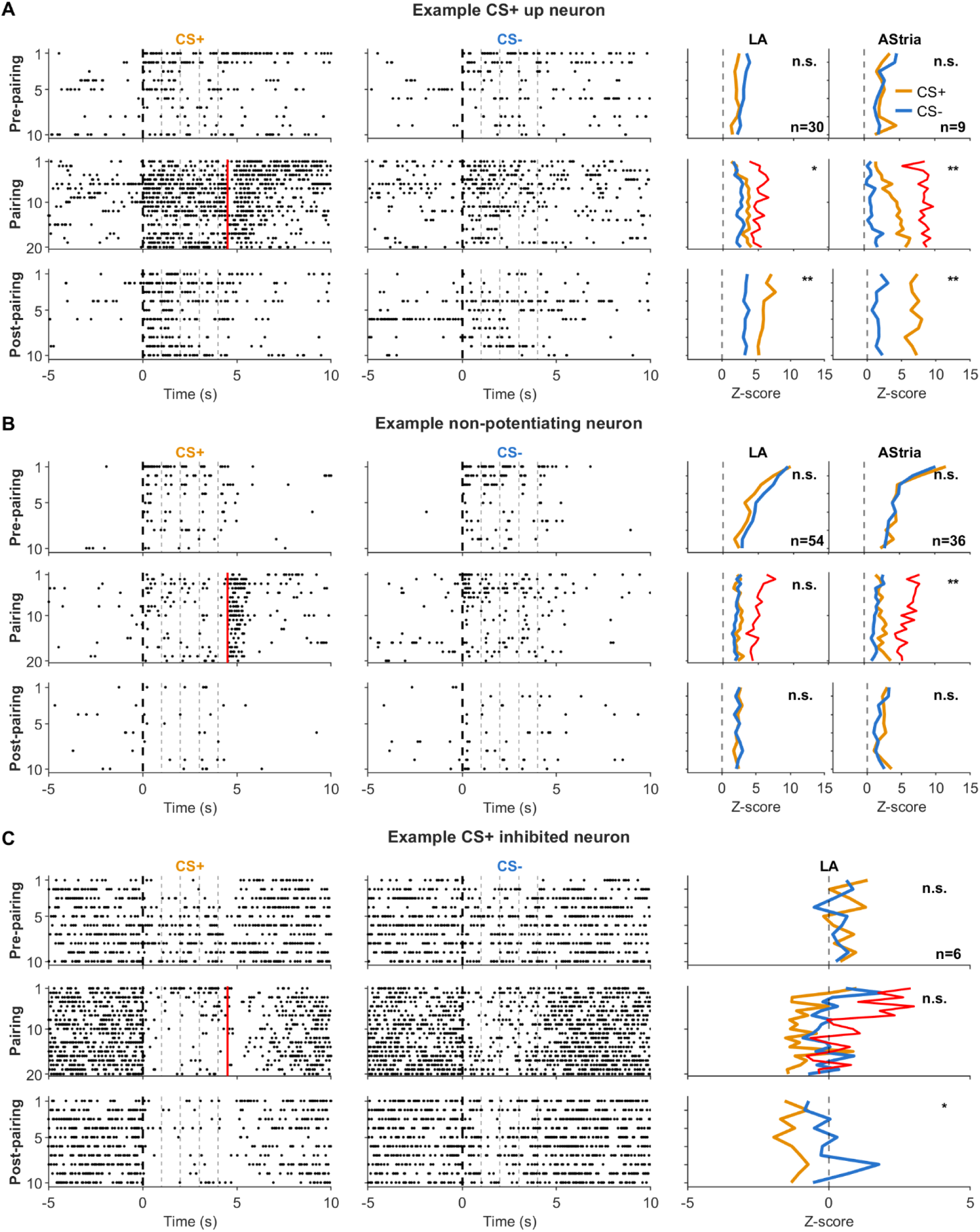
Trial-by-trial z-score dynamics across the three experimental sessions for distinct plasticity classes. Each panel shows data for one example neuron group. Columns 1–2: representative raster plots (CS+ and CS− separately) across pre-pairing, pairing, and post-pairing sessions; each row is one trial. A solid red vertical line at t = 4.5 s marks US onset during the pairing. Column 3: mean per-trial z-score lineplots across neurons in each group (CS+ in orange, CS− in blue; during the pairing session, the US-window response is additionally plotted in red). Wilcoxon signed-rank test was used to compare CS+ versus CS− responses within each session and group. **(A)** CS+ up neurons (CS+↑ class; LA: n = 30, AStria: n = 9). **(B)** Non-potentiating neurons (pooled CS+↓ and CS−↓; LA: n = 54, AStria: n = 36). **(C)** CS+ down neurons that were inhibited (CS+↓ inhibited class; LA: n = 6). For results see Table S15.

In addition to CS+ up neurons, we identified a population of neurons that also exhibited marked habituation during the pre-pairing session and were excited by the US but failed to show increased firing during pairing (Figure 8B). These “non-potentiating” neurons included both CS+ down and CS− down neurons (Figures 7C, D and 8B; Table S15). These findings further indicate that LA and AStria neurons exhibit similar firing dynamics during CS–US pairing, regardless of whether their activity is ultimately up- or down-regulated.

In addition to CS–US pairing-induced potentiation, a previous electrophysiological study identified amygdala neurons that became inhibited during CS presentations following fear conditioning (Lee et al., 2021). Motivated by these findings, we identified several examples of such CS+-inhibited neurons among the CS+ down population in the LA. These neurons exhibited either no response or weak inhibition during the initial CS+ and CS− presentations in the pre-pairing session. When inhibitory responses were present, they habituated over the course of repeated stimulus presentations. During pairing, however, these neurons either regained inhibitory responses or developed inhibition selectively to the CS+, which persisted into the post-pairing session (Figure 8C, Table S15). Thus, LA neurons may acquire not only excitatory but also inhibitory responses as a consequence of CS–US pairing. Such bidirectional plasticity may contribute to enhanced discrimination between the CS+ and CS−.

To determine whether any firing features could predict whether a neuron would undergo pairing-induced up- or down-regulation, we compared baseline firing rate, burst index, and responsiveness to the US in both LA and AStria neurons. None of these measures differed between the groups (Figure S9, Table S16). Therefore, neither intrinsic firing properties prior to pairing nor responsiveness to the US predicted the direction of subsequent plastic changes.

### Spiking dynamics during CS-US pairing

Finally, we examined how responses to the CS+ and US evolved during pairing in CS+ up neurons (Figure 9). Based on their responses to the US, these neurons could be divided into two groups.

**Figure 9.**
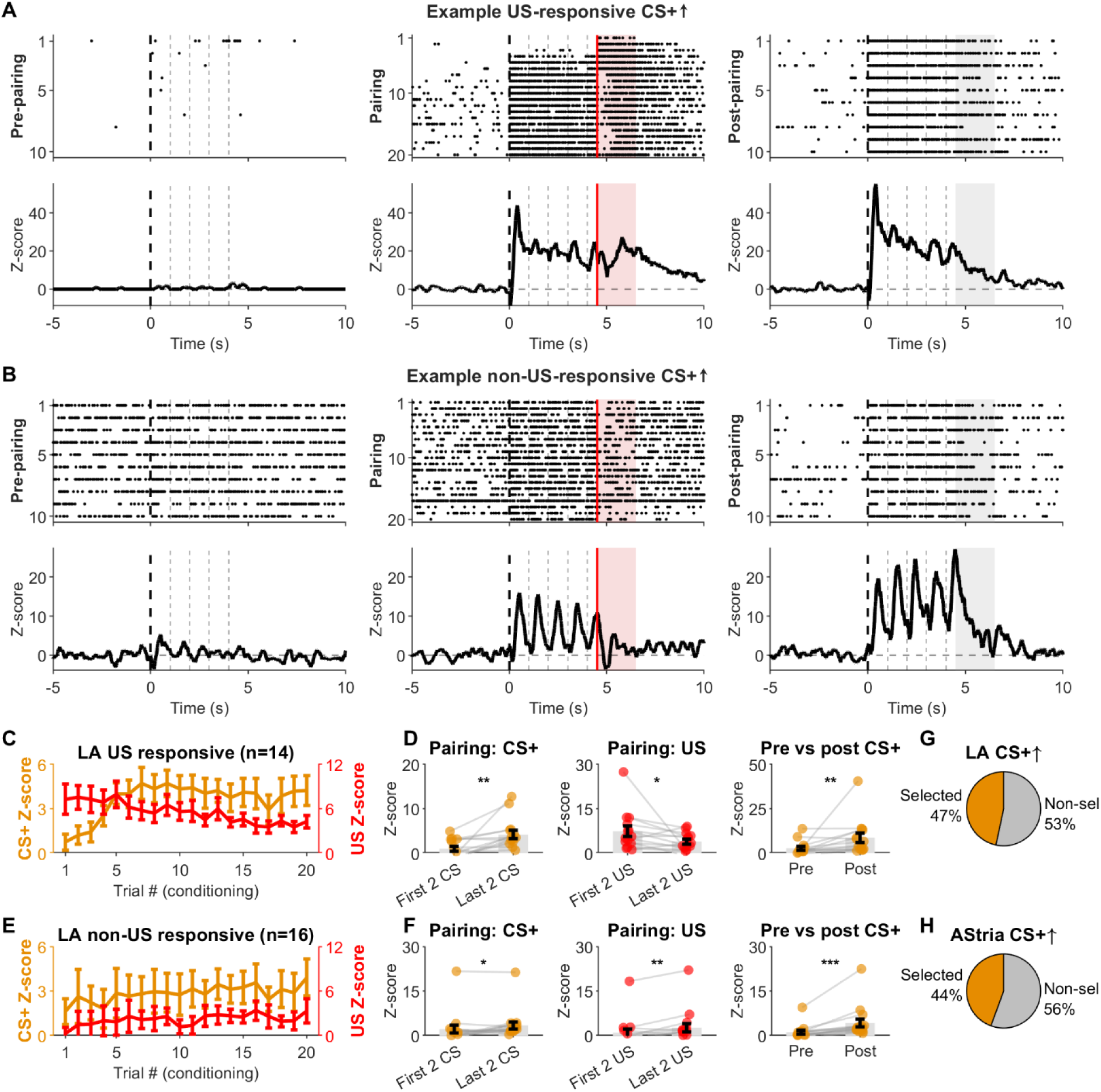
Spiking dynamics in CS+ up neurons across the three experimental sessions. **(A, B)** Representative CS+ up example neurons. Top row: raster plots for CS+ trials across pre-pairing, pairing, and post-pairing. Bottom row: corresponding trial-averaged z-score peristimulus time histograms. In the pairing session, red shading marks the US period (4.5–6.5 s) and a solid red vertical line at t = 4.5 s indicates US onset. In the post-pairing panels, gray shading marks the same time window. (A) US-responsive CS+ up neuron. (B) Non-US-responsive CS+ up neuron. **(C)** Population mean ± SEM per-trial z-score during CS-US pairing for LA US-responsive CS+ up neurons. CS+ z-score is plotted on the left axis (orange); US z-score on the right axis (red). Each point represents one pairing trial. **(D)** Bar graphs for LA US-responsive CS+ up neurons comparing early (first 2 trials) versus late (last 2 trials) z-scores in three windows: CS+ window during pairing (0–4.5 s), US window during pairing, and mean CS+ z-score in pre-pairing (habituation) versus post-pairing (recall). Individual neuron values are overlaid. Wilcoxon signed-rank test. For results see Table S17. **(E)** Population mean ± SEM per-trial z-score during the conditioning phase for LA non-US-responsive CS+ up neurons, formatted as in (C). US z-score uses fixed window (4.5–5.5 s). **(F)** Bar graphs for LA non-US-responsive CS+ up neurons, formatted as in (D). For results see Table S17. **(G, H)** Pie charts showing the proportion of US-responsive versus non-US-responsive CS+ up neurons in LA (G) and AStria (H).

Approximately 47 % of CS+ up neurons in the LA exhibited strong excitation in response to the US at the beginning of pairing, which gradually declined over subsequent pairings (US-responsive neurons). In these neurons, the CS+ initially evoked only a modest increase in firing, but this response progressively strengthened during pairing and reached a plateau after approximately 6–7 pairings (Figure 9A, C, D; Table S17). In contrast, 53% of CS+ up neurons showed little or no response to the US at the onset of pairing (non-US-responsive neurons). Although US responses increased slightly but significantly by the end of the pairing session, they remained relatively weak. Nevertheless, these neurons displayed clear CS+-evoked firing that also increased during pairing (Figure 9B, E, F; Table S17). Importantly, the magnitude of pairing-induced plasticity, assessed by comparing post-pairing and pre-pairing responses, was similar in both groups (US-responsive: 5.94±2.82, n = 14; non-US-responsive: 3.06±0.78, n = 16; Wilcoxon rank sum test, p=0.85; Figure 9D, F). Although fewer CS+ up neurons were recorded in the AStria (n = 9), both the proportion of US-responsive versus non-US-responsive neurons and their firing dynamics during pairing closely resembled those observed in the LA (Figure 9D, F and Figure S10; Table S17). These results indicate that CS+ up neurons can receive either subthreshold or suprathreshold US-driven excitation while developing comparable levels of pairing-induced plasticity.

Overall, our detailed analyses of neuronal activity across the habituation, pairing, and recall sessions strongly support the conclusion that LA and AStria neurons not only respond similarly to sensory stimuli but also exhibit highly comparable response patterns and firing dynamics associated with CS–US pairing.

## Discussion

Our results reveal a previously unrecognized parallel operation between LA and AStria circuits driven by sensory stimuli. Across conditions, tone and shock evoked closely matched firing increase and dynamics in LA and AStria neurons. In contrast, BA and CeA neurons showed little tone-evoked excitation and were dominated by inhibitory responses, indicating that these downstream amygdala nuclei do not mirror the early sensory computations occurring in LA and AStria. Moreover, CS–US pairing induced comparable discriminative plasticity in both regions. These findings suggest that LA and AStria may function as a unit that receives and processes tone and shock signals in a parallel and highly similar manner.

### Parallel sensory processing in LA and AStria

The parallel spike dynamics observed in LA and AStria align closely with their shared anatomical inputs from the same thalamic and cortical areas (LeDoux et al., 1991; Romanski and LeDoux, 1993; Agster et al., 2016; Tsukano et al., 2019; Gehrlach et al., 2020). These shared afferents provide a structural explanation for the nearly identical tone- and shock-triggered spike responses we observed. In particular, the short-latency (<25 ms) responses in both structures likely reflect direct thalamo→amygdala/striatal drive, consistent with previous reports of MGm/PIL→LA conduction delays (Bordi and LeDoux, 1992; Quirk et al., 1995; Li et al., 1996; Barsy et al., 2020). The subsequent activity phase (≈25–35 ms) likely reflects a convergence of cortical afferents and local LA outputs (Stefanacci et al., 1992; Li et al., 1996; Quirk et al., 1997; Abatis et al., 2024; Reeb et al., 2025). More prolonged activity (>35 ms) is likely to rely on recurrent LA microcircuits and reciprocal LA-cortical loops (Lamme and Roelfsema, 2000; Yu et al., 2004; Feigin et al., 2021; Asokan et al., 2023; Abatis et al., 2024). This sequential evolution of neuronal activity within LA and AStria circuits provides a framework for multi-timescale integration of tone and shock signals and is consistent with the prevailing model of information processing in the LA.

### Shock-driven suppression of tone responses

Two observations in our study support the view that tone and shock inputs do not simply summate. First, tone+shock responses were only marginally larger than shock responses alone at longer timescales, indicating that shock-evoked activity dominates the sensory response profile. In addition, no evidence for supralinear summation was found in case of monosynaptically evoked spiking, which is in contrasts with those observed in the MGm/PIL (Barsy et al., 2020). Second, tone-responsive neurons often failed to fire when the tone was co-presented with the shock, revealing a shock-driven suppressive mechanism. The source of this inhibition remains unresolved, but two candidate mechanisms emerge. 1) The shock robustly recruits the parabrachial nucleus, which in turn can excite GABAergic neurons in zona incerta and/or thalamic reticular nucleus (Shammah-Lagnado et al., 1985; Urbain and Deschenes, 2007; Aizenberg et al., 2019; Kirouac et al., 2022; Liu et al., 2022). These GABAergic neurons may suppress spiking of MGm/PIL neurons in a feedforward manner, reducing tone-driven firing in the LA and AStria (Barsy et al., 2020; Taylor et al., 2021). 2) At the local circuit level, the shock may powerfully recruit feed-forward inhibition in LA and AStria, sufficiently strong to override tone-evoked excitation (Windels et al., 2010; Windels et al., 2016). Future studies will be needed to distinguish between these possibilities.

### PV IN-mediated gain control is conserved in LA and AStria circuits

Our study demonstrates that PV INs exert tight feedforward control over the earliest sensory stimulus-evoked spikes in both LA and AStria. Although previous in vivo work showed that PV IN inhibition increases PN firing in the basolateral amygdala during sensory stimulation (Wolff et al., 2014), these studies did not isolate the short-latency monosynaptic window. Prior findings strongly support the role of PV INs in the earliest sensory stimulus-evoked responses in LA and AStria: 1) MGm/PIL afferents form synapses onto PV INs in the LA (Woodson et al., 2000) and 2) Thalamic inputs drive PV IN activity in vitro (Lucas et al., 2016) and in vivo (Barsy et al., 2020). Our data extend these findings by demonstrating that shock-evoked spiking in the AStria is also under PV IN control, indicating that the same feedforward motif is conserved across LA and AStria circuits.

### Sensory stimulation primarily excites BA inhibitory cells

Although tone presentations evoked little excitation in the BA or CeA, shock-evoked activity was readily detectable in the BA. Nevertheless, inhibitory responses predominated within the BA neuronal population. This prevalence of inhibition may be explained, at least in part, by the preferential recruitment of inhibitory interneurons, which exhibited particularly robust increases in firing in response to sensory stimuli. Our findings are consistent with previous studies suggesting that LA activity recruits only a sparse subset of BA neurons (Reijmers et al., 2007; Rashid et al., 2016; Abdou et al., 2018; Abatis et al., 2024). Such neurons may have been underrepresented in our recordings, thereby contributing to the relatively low incidence of excitatory responses observed in the BA. Similarly, shock-evoked excitation in the CeA was generally weak. One possible explanation is that our recordings undersampled neuronal populations that receive strong nociceptive input from the parabrachial nucleus (Han et al., 2015; Sugimura et al., 2016; Chen et al., 2018). Together, these findings suggest that excitatory responses to aversive stimuli in both the BA and CeA may be restricted to relatively small and specialized neuronal populations.

### Similar discriminative plasticity and spiking dynamics in LA and AStria neurons associated with CS-US pairing

In line with prior studies (Grewe et al., 2017; Lee et al., 2021), we observed that CS–US pairing induced four distinct types of changes in spiking activity both in LA and AStria neurons. These findings strongly suggest that pairing-induced plasticity in spiking activity is, at least in part, regulated by upstream brain regions, as the local circuits of the LA and AStria differ substantially in their organization. Consistent with this interpretation, comparable changes in neuronal activity have been reported in the MGm/PIL following CS–US pairing (Taylor et al., 2021). Our experiments further revealed that CS–US pairing can induce not only elevation but also inhibition in spiking, a form of plasticity that is difficult to detect using Ca^2^⁺ imaging. Such bidirectional changes in firing activity may represent an important mechanism underlying engram formation. Finally, we found that 6–7 CS–US pairings were sufficient to produce a maximal increase in spiking activity in CS+ up and CS+ inhibited neurons, closely matching the number of pairings typically used in fear-conditioning paradigms (Wotjak, 2019). Notably, a subset of CS+ up neurons was not strongly excited by the US early during pairing, nevertheless, these neurons likely received subthreshold input, as their US-evoked spiking gradually increased over the course of pairing, as it has been observed earlier (Grewe et al., 2017). These observations suggest that at least two distinct cellular and/or synaptic mechanisms contribute to pairing-induced discriminative plasticity in CS+ up neurons.

### Distributed defensive circuits centered on the LA

Our findings integrate into an emerging framework in which defensive behavior and fear learning are supported by distributed amygdalo-cortico-striatal loops rather than by a hierarchical chain comprised by amygdala nuclei. When LA is placed at the center of this connectome, three major output streams can be identified: 1) Ventral route composed of LA → BA → CeA → PAG/hypothalamus (Johansen et al., 2011) and LA → BA → CeA → substantia nigra pars lateralis (SNL) pathways (Steinberg et al., 2020), 2) Medial route formed by LA → AStria → SNL pathway (Stefanacci et al., 1992; Jolkkonen et al., 2001), and 3) Dorsal route comprised reciprocal LA ↔ cortical loops (LeDoux et al., 1991; Pikkarainen and Pitkanen, 2001; Tsukano et al., 2019; Gehrlach et al., 2020; Valjent and Gangarossa, 2021). Both CeA and AStria converge onto the SNL, which contains aversive stimulus-responsive dopamine neurons (Menegas et al., 2018; Cox and Witten, 2019). These dopamine neurons project back to the AStria (Menegas et al., 2018) and are well positioned to regulate synaptic plasticity in the circuits that process threatening stimuli (Shen et al., 2008; Cox and Witten, 2019). Crucially, MGm/PIL afferents carrying tone and shock signals target all nodes of the dorsal and medial routes (Romanski and LeDoux, 1993; Linke and Schwegler, 2000; Barsy et al., 2020; Cai et al., 2024), providing a unified subcortical drive for distributed neuronal processing during threatening conditions. This complex neuronal network centered on the LA is involved in diverse brain functions, including fear memory formation and storage, cue and context discrimination, controlling passive and active defensive behaviors.

## Conclusion

Although our acute recordings did not allow us to monitor neuronal activity across days, they enabled high-quality characterization of sensory stimulus-evoked spiking activity, response types, and firing dynamics. These results demonstrate that LA and AStria networks process tone and shock signals in parallel, are regulated by PV interneurons, and undergo similar forms of discriminative plasticity following CS–US pairing. This proposed functional unit may constitute a key hub for sensory stimulus processing linked to aversive situations and, thereby, can be an important regulator of defensive behavior and fear learning under physiological and pathological conditions.

## ACKNOWLEDGEMENTS

This work was supported by NIH R21 NS140988, P30 DA054610 and National Research, Development and Innovation Office (K131893) grants. In addition, this study received financial support from the Indiana University Bloomington (IN), Linda and Jack Gill Foundation of Texas (TX), IU Research and HUN-REN Hungarian Research Network. The authors are grateful to Becca Daye for her excellent technical assistance. We also thank Bence Barabás, András Kun and Light Microscopy Imaging Center (LMIC) at the Indiana University Bloomington for kindly providing microscopy support.

